# The limits to parapatric speciation 3: Evolution of strong reproductive isolation in presence of gene flow despite limited ecological differentiation

**DOI:** 10.1101/2020.02.19.956292

**Authors:** Alexandre Blanckaert, Claudia Bank, Joachim Hermisson

## Abstract

Gene flow tends to impede the accumulation of genetic divergence. Here, we determine the limits for the evolution of postzygotic reproductive isolation in a model of two populations that are connected by gene flow. We consider two selective mechanisms for the creation and maintenance of a genetic barrier: local adaptation leads to divergence among incipient species due to selection against migrants, and Dobzhansky-Muller incompatibilities (DMIs) reinforce the genetic barrier through selection against hybrids. In particular, we are interested in the maximum strength of the barrier under a limited amount of local adaptation, a challenge that may initially face many incipient species. We first confirm that with classical two-locus DMIs, the maximum amount of local adaptation is indeed a limit to the strength of a genetic barrier. However, with three or more loci and cryptic epistasis, this limit holds no longer. In particular, we identify a minimal configuration of three epistatically interacting mutations that is sufficient to confer strong reproductive isolation.

## Introduction

Understanding the mechanisms that drive speciation remains a challenge of evolutionary research [1, 2, 3, 4]. Recently, parapatric speciation - where incipient species are spatially separated, but still exchange migrants - has received considerable attention, both in empirical and theoretical research [5, 3, 6, 7]. In particular, several studies have analysed the potential for the evolution of a postzygotic isolation barrier in the presence of gene flow [5, 8, 9]. Whereas such barriers can easily arise in strict allopatry, even small amounts of gene flow can impede their buildup. This is due to two main problems. First, persistent gene flow acts to swamp divergent alleles between populations [5]. Second, gene flow creates a permanent fitness cost for any genetic incompatibility that contributes to a genetic barrier due to production of unfit hybrids [9]. Local adaptation can be a potent mechanism to protect divergent alleles from swamping. Indeed, there are indications that at least some local adaptation is necessary for parapatric speciation [10, 5]. Consequently, some authors [11] have suggested mechanisms purely based on divergent selection to explain how speciation can happen in parapatry. They assumed that each new mutation contributes to local adaptation. Barrier genes are additive without epistasis between single mutations. This corresponds to a scenario of pure ecological speciation. With such unlimited potential for ecological differentiation, evolution can easily build a genetic barrier to gene flow. However, this is not necessarily a realistic mechanism in natural populations. Whereas immigrants from a genetically closely related sister population may often have fitness deficits, they are rarely “dead on arrival”. Especially early during divergence, environments need to be similar enough for the ancestral population to survive in both habitats. This limits the selection differential generated by local adaptation. For example, Via [12] showed in pea aphids that residents have a fitness that is 3.3 to 20 times larger than the fitness of migrants. Furthermore, genetic barriers that are based uniquely on ecological differences can only be temporary, since they are maintained only as long as their causal environment persists. The dissociation between local adaptation and the strength of a genetic barrier to gene flow is thus key for the evolution of strong reproductive isolation and for completing the speciation process.

In this manuscript, we address when and how strong reproductive isolation can evolve between two parapatric populations with limited ecological differentiation. To this end, we first define measures that characterize the strength of a genetic barrier and compare this with the amount of ecological differentiation that is available between the two populations. We then focus on the role of epistasis and the pattern of incompatibilities among genes building the genetic barrier. Our results show that, for a broad range of conditions, the potential for ecological differentiation is indeed an upper limit for the strength genetic barrier that can be formed. However, we also show that this constraint can be broken and that particular patterns of strong epistasis enable the evolution of strong reproductive isolation in parapatry. A barrier of this type must involve at least three interacting loci: two interacting barrier loci and one locus that changes their genetic background. A strong genetic barrier can thus evolve parapatrically in (minimally) three steps from an undifferentiated initial state.

## Model

### General definition

We consider a migration-selection model in a continent-island framework [5, 9]. The model consists of two panmictic populations, one on an island and the other on a continent, each of sufficient size that we can ignore genetic drift. We consider the population genetic dynamics of the island population, which receives unidirectional migration from the continental population at (backward) rate *m* per individual and generation. In the main part of this article, we consider a three-locus model, with diallelic loci *A, B* and *C*. Ancestral alleles are denoted by lowercase letters and derived alleles by uppercase letters. Allele *A* (resp. *B* and *C*) has a selection coefficient *α* (resp. *β* and *γ*) compared to the ancestral allele *a* (resp. *b* and *c*). We derive extended results for models with more than three loci in the Supplementary Information (SI). Below and in the SI, for multiple loci *A*_*i*_ and *B*_*j*_, we use the following notation: allele *A*_*i*_ has a selective advantage *α*_*i*_ over allele *a*_*i*_ and its epistatic interaction with allele *B*_*j*_ is given by 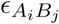.

Epistasis can occur between any combination of derived alleles and is denoted by *ϵ*, with the involved alleles indicated as subscript. For example, *ϵ*_*ABC*_ denotes 3-way epistasis between alleles *A, B* and *C*. The fitness of each haplotype is given in Table 1.

**Table 1:** Notation of frequencies *x*_*i*_ and fitness values *w*_*i*_ of the eight different haplotypes for haploid populations in the 3-locus model.

| Hap. | abc | Abc | aBc | abC | ABc | AbC | aBC | ABC |
| --- | --- | --- | --- | --- | --- | --- | --- | --- |
| $x_i$ | $x_1$ | $x_2$ | $x_3$ | $x_4$ | $x_5$ | $x_6$ | $x_7$ | $x_8$ |
| $w_i$ | 0 | $\alpha$ | $\beta$ | $\gamma$ | $\alpha + \beta$ | $\alpha + \gamma$ | $\beta + \gamma$ | $\alpha + \beta + \gamma$ |
| | | | | | $+ \epsilon_{AB}$ | $+ \epsilon_{AC}$ | $+ \epsilon_{BC}$ | $+ \epsilon_{AB} + \epsilon_{AC} + \epsilon_{BC} + \epsilon_{ABC}$ |

We assume that the continent is always monomorphic. When evolution happens on the continent, each substitution is assumed to be instantaneous. That is because we are only interested in the (potential) polymorphic equilibrium state on the island, where individuals from both populations meet and mix. The haplotype frequency dynamics of an arbitrary haplotype *X* on the island (e.g., *X* = *abC*) under the continuous-time weak-selection approximation is:

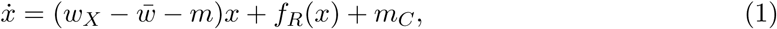

where the migration rate *m*_*C*_ = *m*, if *X* is the continental haplotype and *m*_*C*_ = 0 otherwise. Here, *f*_*R*_(*X*) describes the change in frequency of haplotype *X* due to recombination. The detailed ordinary differential equation system with an explicit expression of the complicated function *f*_*R*_(*X*) is given in the SI (eq. S1). Our analytical results focus on two special cases that have been shown to capture most of the important behaviour [5, 9]: tight linkage (defined as the limit *r →* 0 for all recombination rates, *f*_*R*_(*x*) = 0) and loose linkage (defined as the limit *r → ∞*; dynamics are given in eq. S4). The second scenario corresponds to the assumption of linkage equilibrium between all loci, which approximately holds true when the recombination rates are much larger than the selection coefficients and migration rates [5, 9] (confirmed in the Results section Fig. 3). We complement our analytical approach with numerical simulations for intermediate recombination rates.

**Figure 1:**
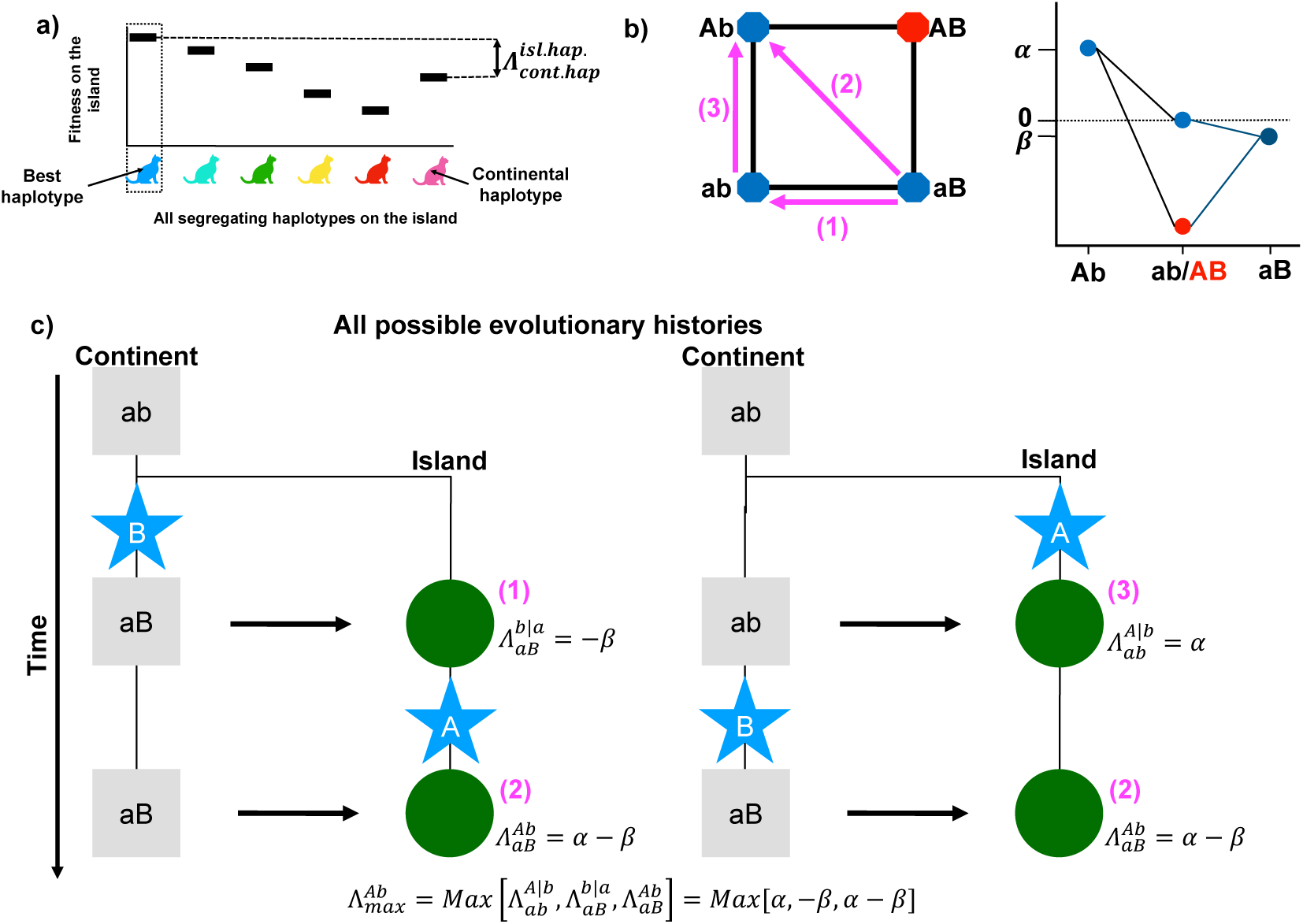
Measures of local adaptation. We define two measures of environmental heterogeneity between the continent and the island, the “current amount of local adaptation” and the “maximum amount of local adaptation”. a) The schematic shows an example in which six haplotypes are segregating on the island. The current amount of local adaptation of the population, 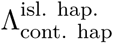 corresponds to the difference in fitness, evaluated on the island, between the fittest segregating possible haplotype on the island (in blue) and the fittest possible haplotype on the continent (in pink). b) Fitness graph and fitness landscapes for a two-locus model with a DMI. The arrow corresponds to the fitness comparison between the continental haplotype (base of the arrow) and island haplotype (tip of the arrow), with the number corresponding to the evolutionary step in panel c). The fitness landscape shows a case in which *β* < 0, meaning that *B* is a local adaptation to the continent. In our general model, *β* can take both positive and negative values, which means that *B* can also be beneficial both on the island and the continent. c) Potential evolutionary histories leading to the formation of a genetic barrier in a 2-locus model. For each possible evolutionary step, we compute the current amount of local adaptation of the island population as 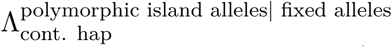. The magenta numbers correspond to the same comparison made on the fitness graph from b). The maximum amount of local adaptation, 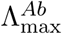, generated by the fitness graph given in panel, is the maximum of these values. If we use the fitness landscape depicted in panel b), we obtain 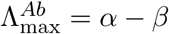.

**Figure 2:**
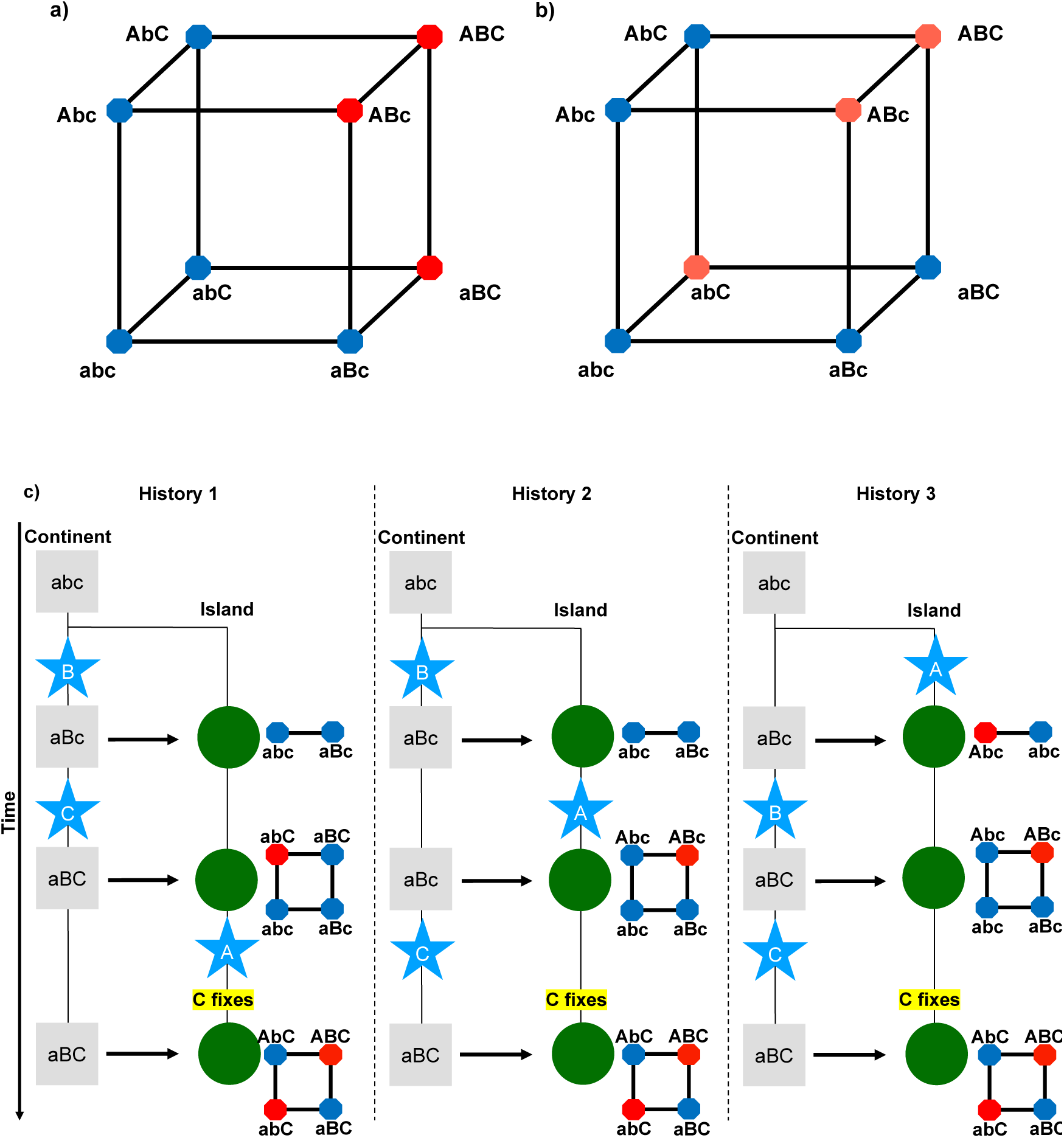
With 3 loci and cryptic epistasis, the high-fitness ridge of a parapatric 2-locus DMI can be turned into a deep fitness valley. a) Fitness graph for a model with negative pairwise epistasis between *A* and *B*, and *B* and *C*, which does not allow for parapatric evolution of a strong genetic barrier. Red dots correspond to low fitness haplotypes and blue dots to high fitness haplotypes. b) Fitness graph for a model with negative epistasis between *A* and *B* and a strongly deleterious allele *C*. Both alleles *A* and *B* can compensate for the deleterious effect of *C* but the compensation is not cumulative. This fitness landscape can allow for the parapatric evolution of a strong genetic barrier, because it contains a 2-locus fitness graph with two fitness peaks isolated from each other by a deep valley, if allele *C* is fixed. c) Three possible evolutionary histories and the temporary underlying fitness graphs (subgraphs of b)) can lead to the formation of a fitness landscape in which the two fitness peaks (*AbC* and *aBC*) are separated by an unsurpassable fitness valley. This strong genetic barrier can evolve via single-step mutations in the presence of gene flow, due to the existence of a high fitness ridge that disappears through fixation of allele *C*.

**Figure 3:**
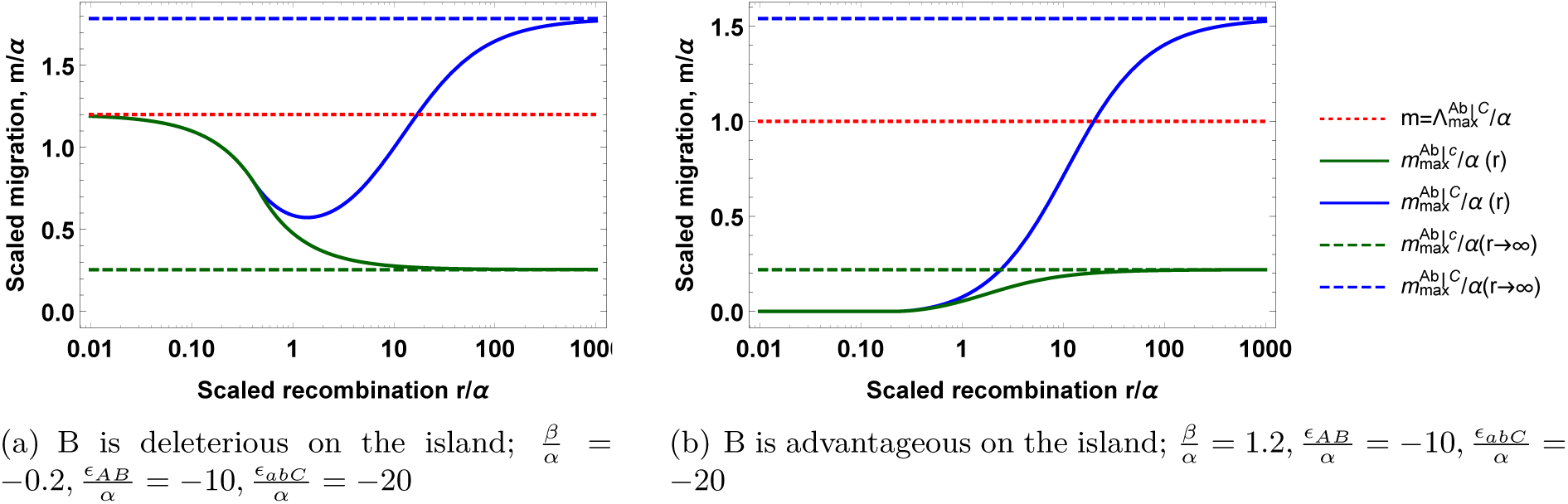
Strong genetic barriers form through global fixation of allele *C* if there is sufficient recombination. The relative maximum amount of local adaptation 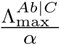 is given by the dotted red line. The graph shows the relative strength of the genetic barrier as a function of recombination for the two-locus DMI before the *C* mutation appears, 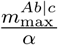 in green, and after *C* has fixed on the island 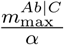 in blue. Their corresponding limits for loose linkage are represented by the dashed lines, in green for 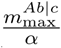 and in blue for 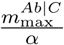. The limits for tight linkage are for panel a) 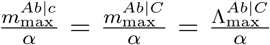 (red dotted line) and for panel b) 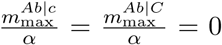. All parameters are scaled by the direct selective advantage of allele *A*.

We study both haploid and diploid populations. For diploid populations, we assume that all direct effects of derived alleles are codominant [5, 9, 13]. Regarding epistasis, we consider two scenarios: codominance and recessivity of the epistatic interaction. The two scenarios differ in the expression of epistasis in double and triple heterozygotes (cf. [5]).

With the continuous time approach employed here, all selection and migration parameters are rates. For the study of equilibria, only relative rates matter and we can thus scale all parameters by the selective advantage of the *A* allele on the island, *α* (note that we always assume *α ≠* 0).

### Measures of the genetic barrier to gene flow and the maximum amount of local adaptation

Our aim here is to assess scenarios in which a strong barrier to gene flow can evolve despite limited potential for (extrinsically driven) local adaptation. To this end, we need to define measures for both the barrier strength and the amount of local adaptation.

Following Bank et al. (2012) [5] and Blanckaert & Hermisson (2018) [9], we define the barrier strength as the maximum migration rate *m*_max_ that can be sustained while maintaining the polymorphism at the barrier loci. Note that *m*_max_, defined this way, is specific to a set of barrier loci in a specific genetic background. We reflect this in our notation by adding labels to *m*_max_ to indicate the island alleles that are maintained polymorphic. For example, consider a 2-locus barrier with derived alleles *A* and *B* at the barrier loci, with allele *A* appearing on the island and *B* on the continent. To maintain both loci polymorphic, alleles *A* and *b* must persist on the island in migration-selection balance, because *aB* is the immigrating haplotype from the continent. 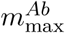 is the maximum migration rate for the maintenance of this stable equilibrium; above this value either *A* or *b* (or both) are lost. The barrier strength can also depend on the genetic background. We will include reference to this background in our notation whenever necessary by writing 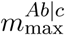 or 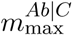 for the strength of the (*Ab*) barrier in the background of the ancestral *c* or derived *C* allele, respectively, where either of these alleles at locus *C* is fixed on both the continent and the island. While others measures exist (e.g., introgression probability of a linked neutral allele [14]), we focus on this measure, which assesses the maintenance of divergence at the barrier itself.

To measure local adaptation, we define two parameters that capture either the current state or the overall fitness landscape of the system. The first one, Λ, depends on a subset of model parameters and the time of observation, the second, Λ_max_, depends on the whole set of model parameters.

For any state of the population, we measure the *current amount of local adaptation* on the island Λ as the fitness advantage of the fittest segregating genotype on the island over a continental migrant. Recall that the continent is always monomorphic in our model. (With a polymorphic continent, the genotype with the largest fitness on the continent would provide the reference). This measure is consistent with the verbal notion of local adaptation by Kawecki & Ebert (2004) [15] and illustrated in Figure 1. Using the 2-locus barrier example mentioned above, the current amount of local adaptation, after the first mutational step, is given by 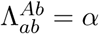 if *A* appeared first and 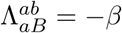 if *B* appeared first. After the second mutational step, the current amount of local adaptation is given by 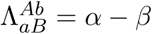.

In addition, we define the *maximum amount of local adaptation* that can occur in the model over the course of the differentiation process that results in a given genetic barrier as Λ_max_. Note that Λ_max_ does not depend on the current state, but is a property of the full fitness landscape. It captures all states that could have occurred (i.e., that are allowed by the fitness landscape) during the adaptive process from a given ancestral state. We thus need to consider all possible evolutionary histories to determine Λ_max_. Using the 2-locus barrier example mentioned above, the maximum amount of local adaptation, 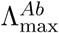 is given by: 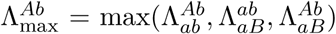. To match the genetic barrier notation, we will use 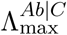 if we need to mention that the genetic barrier depends on the genetic background (here a fixed allele *C*).

From this definition we see, in particular, that the maximum amount of local adaptation for a large barrier which includes many loci is always larger or equal than the value of Λ_max_ for any smaller barrier that involves only a subset of these loci. For diploids, we consider the fitness differences between genotypes scaled by the ploidy of the individual. Using this definition allows us to maintain a consistent notation for haploid and diploid populations: for a single locus *A*, we always have 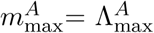. We include a limit to local adaptation into our model by assuming that Λ_max_ is bounded by the ecology of the system. However, the fitness difference between the optimal island genotype and a hybrid (or any maladapted genotype) may be much larger, since these genotypes are not part of any evolutionary trajectories.

## Results

### Maximum amount of local adaptation as a limit to barrier strength

If the external environment sets a limit to the amount of local adaptation, does this also imply a limit on the strength of the genetic barrier that can evolve in the presence of gene flow? We address this question by asking whether the former restricts the latter, i.e. whether the maximum amount of local adaptation during the differentiation process Λ_max_ limits the barrier strength *m*_max_. For simplicity, we will refer to genetic barriers as strong if *m*_max_ > Λ_max_ and as weak otherwise. Indeed, we find that for many types of fitness landscapes and linkage architectures, genetic barriers can only be weak in this sense.

For a single-locus barrier in a haploid population, it is straightforward to see that *m*_max_ = Λ_max_ since local adaptation (direct selection against migrants) is the only mechanism that can maintain a polymorphism. This result holds independently of whether a locally adaptive allele appears on the island or whether a maladaptive allele immigrates from the continent. This result readily generalizes to the case of *n* biallelic loci in tight linkage, which acts like a single-locus model with 2*n* alleles. Only two haplotypes can be maintained at equilibrium [16]: the best one on the island (verifying eq. (S11)) and the immigrating one, regardless of epistasis. This result extends to diploid individuals as long as there is no under- or overdominance. If gene flow exceeds the temporary amount of local adaptation, *m* > Λ, the continental haplotype replaces the island haplotype. Since Λ ≤ Λ_max_, the maximum amount of local adaptation Λ_max_ is always an upper bound for the strength of the genetic barrier, *m*_max_ ≤ Λ_max_. For weak, but non-zero recombination *r*, this result remains valid as long as *r* is small relative to the selection coefficients and migration rates (Fig. S7).

In the absence of epistasis and for multiple loosely linked loci, the temporary amount of local adaptation Λ is simply the sum of the selection coefficients *α*_*i*_ > 0 of segregating island alleles relative to the immigrating continental alleles at the same locus (where island alleles can be ancestral or derived). During the differentiation process, this value is maximized in the final state when all mutations that contribute to the barrier under consideration have appeared, 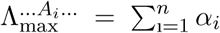. In contrast, the strength of the genetic barrier maintaining all loci *A*_*i*_ polymorphic is given by the smallest selection coefficient: 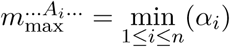. For given Λ_max_, this barrier is therefore maximized when all loci share the same selection strength: 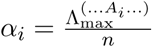. Clearly, we have *m*_max_ < Λ_max_ for more than a single locus, i.e., the maximum amount of local adaptation Λ_max_ is again an upper bound for the strength of the genetic barrier. This result (*m*_max_ < Λ_max_) readily extends to intermediate recombination rates as recombination ends up breaking the best haplotype (once formed) without any additional benefits.

Having shown that it is impossible to form a strong genetic barrier in the absence of epistasis or if all loci are in tight linkage, we now turn to the case with loose linkage and epistasis. This introduces the possibility of selection against recombinant hybrids. Since fitness differences between the optimal types and maladaptive hybrids can be much larger than the strength of local adaptation Λ_max_, selection is not constrained by the ecology and can potentially result in a strong barrier. For two loci and negative epistasis (i.e., a DMI), the barrier strength under a combination of local adaptation and selection against hybrids has previously been analyzed by Bank et al. [5]. The authors focused on the case of an allele *A* appearing on the island and *B* appearing on the continent, with negative epistasis between the two derived alleles. From their result for 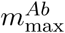 (eq. 11 of [5]), we can deduce that the maximum strength for the corresponding genetic barrier (eq. (2))

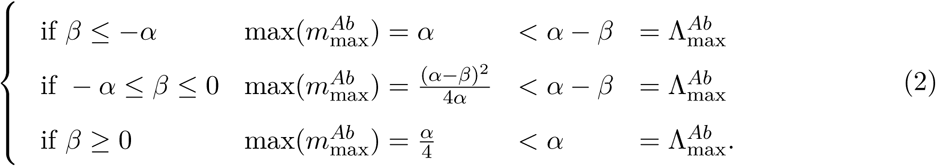

From equation (2) it is clear that the maximum amount of local adaptation is again an upper bound for the strength of the genetic barrier.

We extended this model to allow for positive epistasis and derived the expression for *m*_max_ (given in eq. (S5)). With positive epistasis, a genetic barrier can exist only if allele *B* is deleterious on the island, and the maximum of this barrier is given by 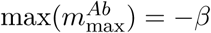. We therefore always obtain *m*_max_ ≤ Λ_max_ = *α − β*. However, in contrast to the negative epistasis case, it is possible for a genetic barrier to reach 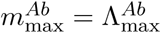, when *A* is neutral (*α* = 0) on the island, and *B* is extremely deleterious 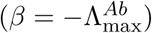 on the island when associated with allele *a* but neutral when associated with allele *A*. This corresponds to a scenario in which allele *A* compensates the deleterious effect of allele *B*. Here, immigration of *B* boosts the marginal fitness of allele *A* and therefore counteracts the swamping effect of immigration of *a*. This result also holds if the roles of *A* and *B* are reversed and if both alleles *A* and *B* appear on the island or on the continent.

For two biallelic loci, there is only a single epistasis parameter. In particular, interactions among derived alleles must be either negative or positive. This severely limits the complexity of the fitness landscape. We identify further, more complex, classes of epistasis patterns, where the maximum amount of local adaptation is an upper bound for the strength of the genetic barrier, as illustrated below for three loci and with general results presented in the SI. These patterns include 1) any barrier that includes either an island allele that is not involved in positive interactions, or a continental allele that is not involved in negative interactions (see sections S 2.3 and S 2.4); 2) any barrier where all derived alleles originate on the island or all on the continent (see sections S 2.5 and S 2.6); 3) any barrier with only positive or only negative epistatic interactions between derived alleles (this directly follows from points 1 and 2) (section S 2.7); 4) any barrier where derived alleles on the continent and the island do not interact, or interact only through negative epistasis (see section S 2.8).

This suggests that only more complex epistasis, with a combination of positive and negative interactions, can result in a strong genetic barrier. We thus consider a diallelic 3-locus model in the rest of the manuscript, which is fully parametrized with three direct selection coefficients and four epistatic parameters, allowing for complex interactions.

### Three-locus model and the role of cryptic epistasis in the formation of strong genetic barriers

#### Haploid populations

We first consider a case of with two pairwise epistatic interactions. First, we focus on a case with two island adaptations *A* and *C*, which appear on the island, and a continental adaptation *B*. The different possible cases illustrate the general rules above for the impossibility of a strong barrier.

- Negative pairwise epistasis between *A* and *B* and *B* and *C* cannot result in a strong barrier. Indeed, in the absence of allele *B*, we have 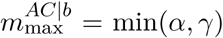, which is smaller than 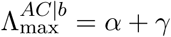. Once allele *B* is introduced on the continent, the marginal fitness of alleles *A* and *C* decreases, leading to 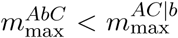. Since the two-locus barrier with *A* and *C* is a subset of the three-locus barrier, 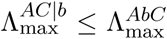, and therefore 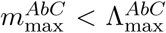. The corresponding fitness graph for this case is given in Fig. 2 panel a).
- Similarly, pairwise positive interaction between *A* and *B*, and *B* and *C* is not sufficient for a strong barrier. The genetic barrier formed by allele *A* and *C*, assuming *B* is fixed on the island, corresponds to a case described above (i.e. two non-interacting loci), therefore 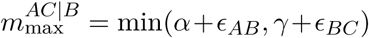, while 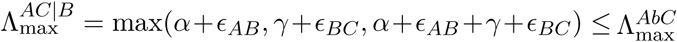. If locus *B* is polymorphic on the island, then the marginal fitness of both allele *A* and allele *C* is reduced, leading to 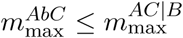 and therefore, 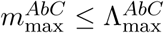.
- We now consider that one pairwise epistatic interaction is positive and the other negative: we assume that alleles *A* and *B* interact negatively and alleles *B* and *C* interact positively. In the absence of allele *C*, this corresponds to the two-locus case mentioned above and therefore 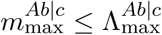. If allele *C* appears on the island, it directly increases the marginal fitness of allele *B* on the island, facilitating its fixation on the island. In addition, through this effect on *B* (leading to a higher equilibrium frequency for *B*), it also indirectly and negatively affects the marginal fitness of allele *A* facilitating its loss. As a consequence of its effect on the marginal fitness on alleles *A* and *B*, we obtain 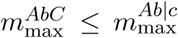 and 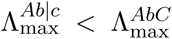, since the “*Ab*|*c*” barrier is a subcase of the “AbC” barrier, leading to 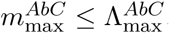.
- Finally, we consider that *A* and *B* interact negatively and *A* and *C* positively. In the absence of allele *B*, the genetic barrier obtained in loose linkage is smaller than its equivalent in tight linkage since recombination breaks the association between *A* and *C*. Or in tight linkage, the genetic barrier is equal to 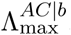. Therefore, in the loose linkage case, 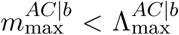. Once the B mutation is introduced, the marginal fitness of allele *A* and *C* decreases due to the direct (for *A*) and indirect (for *C*) interaction with allele *B*. We therefore obtain 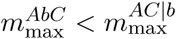; a strong genetic barrier is therefore impossible.

Similar arguments show that any barrier must be weak for two derived barrier alleles on the continent and one on the island (see section S 1.3). The fitness landscapes of all scenarios described so far share a crucial property (Fig. 2a) and S4): the continental haplotype and the fittest island haplotype are connected by a fitness ridge (e.g. *AbC* and *aBc* in Fig. 2a)), and all genotypes on this fitness ridge can be reconstructed from the (fittest) island and continental haplotypes by recombination (recombination of. *AbC* and *aBc* in Fig. 2a)).

We now consider a case in which a genetic barrier with two barrier loci is combined with a change in the genetic background (through a derived allele at a third locus that fixed on both the continent and the island). We assume (as above) that there is an incompatibility between an adaptation on the island at locus *A* and a continental adaptation at locus *B* (i.e. *α* > 0, *ϵ*_*AB*_ < 0 and *β* < −*ϵ*_*AB*_). In addition, we assume that a mutation can occur at a third locus (the *C* locus). We assume that the derived allele *C* is deleterious in the ancestral genetic background (*γ* < 0), but beneficial in the presence of either the *A* or the *B* allele (*ϵ*_*AC*_ > 0, *ϵ*_*BC*_ > 0 and *ϵ*_*ABC*_ ≤ 0; below we assume *ϵ*_*AC*_ = *ϵ*_*BC*_ = −*ϵ*_*ABC*_). If *C* originates on the continent, it can fix on both the continent and the island (eq. (S21)-(S28); see Fig. 2c) for the three potential evolutionary histories). We then obtain a 2-locus barrier (loci *A* and *B*), but the derived alleles at this barrier interact with a fixed derived allele in its genetic background. We refer to this type of interaction as “cryptic epsitasis” since it will not be detected in a study that focuses on divergent alleles between both populations. Notably, the corresponding fitness graph, illustrated in Fig. 2c) (last row), is characterized by the existence of two haplotypes (*AbC* and *aBC*) whose recombinants (*abC* and *ABC*) have very low fitness. Fixation of *C* thus deepens the observed fitness valley between *Ab* and *aB*.

To simplify the notation, we define *γ*′ as the effect of the mutation *C* in the background of at least one other derived allele: *γ*′ = *γ* + *ϵ*_*AC*_. Notably, this system is equivalent to a *C* mutation that appears on the continent, which is advantageous on the island while generating strong negative epistasis with the ancestral background *ab, ϵ*_*abC*_ = −*ϵ*_*AC*_. For the rest of the manuscript, we will use the alternative notation (*ϵ*_*abC*_ and *γ*′) as it is more convenient.

For a haploid population and loose linkage, the dynamics simplify to the classical 2-locus model [5] and are therefore identical to the diploid model (up to some reparametrization, eq. (S14)). The expression for the maximum amount of local adaptation, generated in this model, is

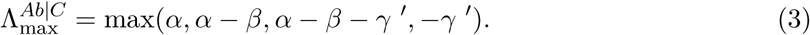

This equation has a simple form because the *abC* haplotype is deleterious and therefore no longer a potential step for evolution. Equation (3) can be reduced to 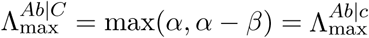 when *C* is advantageous (*γ*′ > 0) on the island (eq. (S19)). The maximum amount of local adaptation, which characterizes the ecological differentiation in the model, is unaffected by the new mutation; *C* modifies the genetic background of both populations but is not directly involved in the divergence process. Since we assume that the new mutation *C* fixes, its position in the genome is irrelevant for the polymorphic equilibrium state. (For conditions of fixation of allele *C* on the island see section S 3.3).

We investigated the impact of this change of the genetic background for two cases analytically: loci *A* and *B* are in tight linkage or in loose linkage. Our analysis was complemented with simulations for intermediate recombination rates (Fig 3). With tight linkage, the barrier remains unchanged in comparison to the original background (at equilibrium, haplotype *abC* does not occur anyway). The barrier is therefore again limited by the maximum amount of local adaptation available, 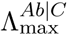. With loose linkage, the genetic barrier can exceed 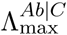 when selection against hybrids, via the strength of epistasis, is strong enough:

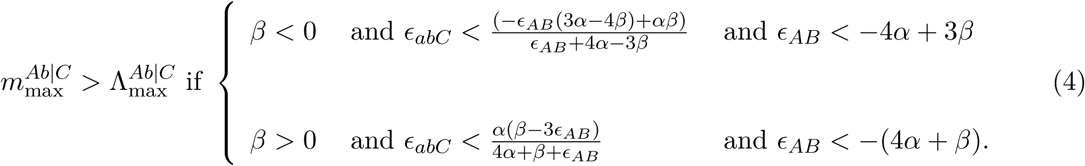

From the previous section, we know that a strong single-locus barrier can never form. However, the existence of a single-locus (unstable) equilibrium is a necessary condition for the 2-locus genetic barrier to be globally stable (SI, section S 3.1.1). Therefore, if 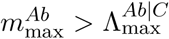, then the 2-locus genetic barrier can only be locally stable, emphasizing the important role of selection against hybrids. When compared to the old barrier 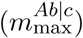, the genetic barrier is strengthened if *-ϵ*_*AB*_ > *α*, i.e, when the incompatibility between *A* and *B* is stronger than the direct selective advantage of *A*; this is therefore a necessary condition for 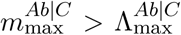. We calculated the genetic barrier numerically for an arbitrary genetic distance between *A* and *B* (Fig. 3): as soon as recombination is strong enough (as selection against hybrid depends both on the formation of those hybrid and their fitness deficit), we recovered the results from loose linkage, independently of the selective advantage of allele *B* on the island. Finally, we also investigated, in the case of loose linkage, the possibility of locus *C* becoming polymorphic instead of being always fixed for allele *C* and showed that strong barriers can also form in these conditions (Fig. S10).

Our assumptions of loose linkage and the continuous-time approximation both implicitly rely on weak selection. We therefore derived the equivalent of 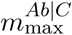 in the discrete-time model assuming that both *abC* and *ABC* are inviable haplotypes and that *A* and *B* are located on different chromosomes. The results are qualitatively similar, i.e. for a range of parameters, a genetic barrier can be stronger than the maximum amount of local adaptation (eq. (S33), Fig. S11). Finally, if we assume that the *abC* haplotype is inviable (*ϵ*_*abC*_ *→ −∞*), the genetic barrier is given by 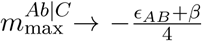. Whereas in the simple 2-locus model before, the barrier strength was limited by local adaptation (which is limited), the formula here shows that the limit is now set by the strength of the incompatibility by hybrid fitness deficit (which is not limited).

#### Diploid populations

In the diploid model we assume that the direct effects of the mutations (*α, β, γ*′) are additive and that epistatic interactions (*ϵ*_*AB*_, *ϵ*_*abC*_) can either be recessive or codominant (see section S 3.2.1). Both the recessive and codominant model simplify to their equivalent dynamics presented in [5], with the same substitutions as in the haploid model (eq.(S14)).

We have established above that the maximum amount of local adaptation, Λ_max_, is not a limit to the strength of a genetic barrier for haploid populations, if epistatic interactions are complex and include interactions with the genetic background. Also in diploids, the strength of the genetic barrier exceeds the maximum amount of local adaptation when negative epistasis is strong enough (area below the line on Fig. 4). More precisely, the maximum amount of local adaptation is not a limit to the strength of the genetic barrier as long as the incompatibilities are strong and expressed in the F1 generation. They can be expressed either through recombination (*A* and *B* in loose linkage) or through the codominance of the interactions. When *A* and *B* are in tight linkage and epistasis is recessive, the genetic barrier is given by 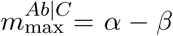 and is therefore at best equal to the maximum amount of local adaptation.

**Figure 4:**
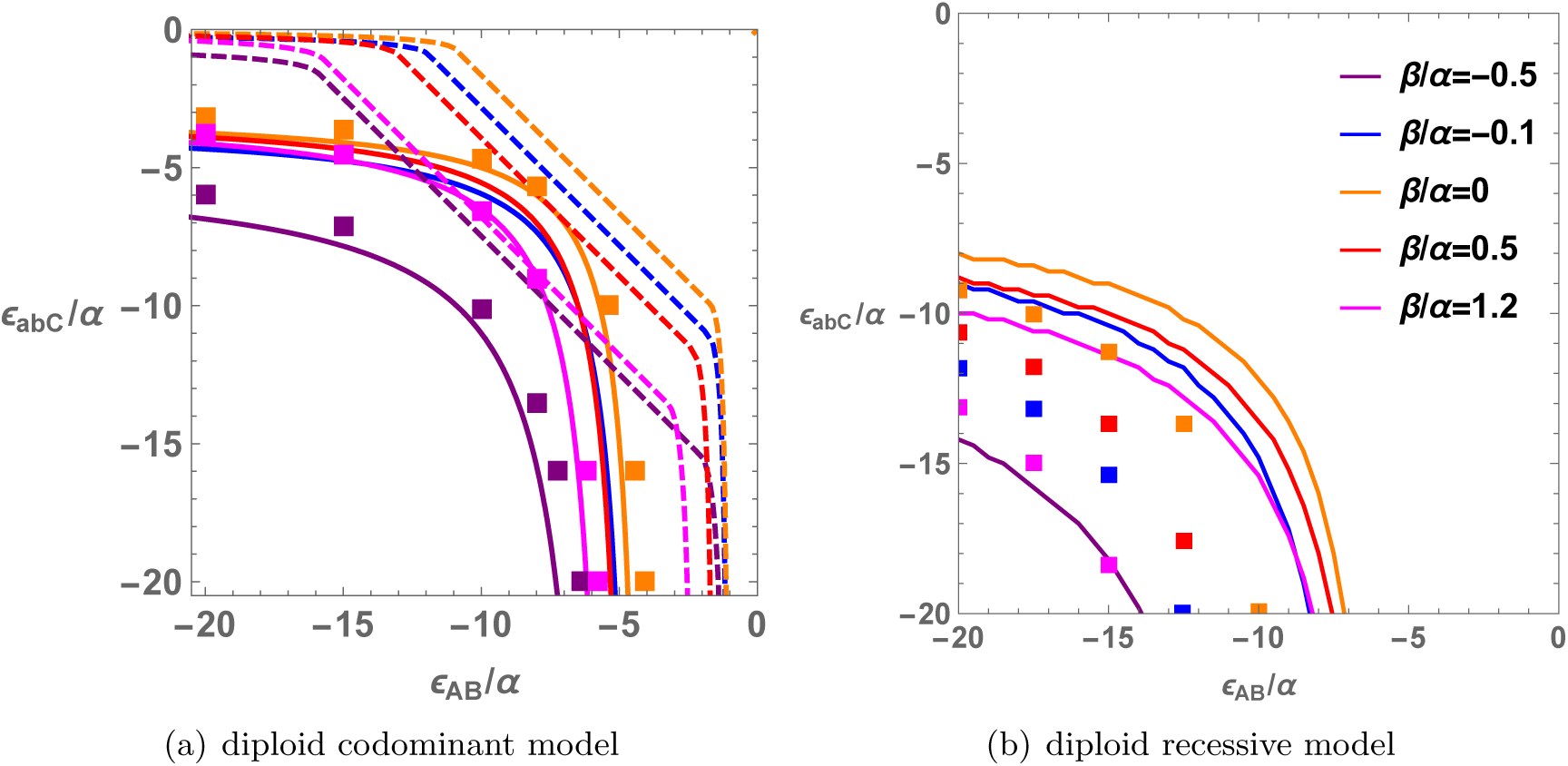
Parameter region for strong genetic barriers in diploids. The X axis corresponds to the epistasis between *A* and *B*, 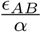. The Y axis corresponds to the epistasis between *C* and the ancestral background, 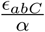. The area below each curve (strong negative epistasis) indicates the parameter region with a strong genetic barrier, 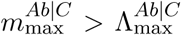. Each color corresponds to a different value of 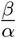, ranging from locally maladapted alleles *B* for 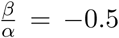 to strongly beneficial alleles *B* on the island for 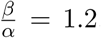. The left panel corresponds to the codominant diploid model, with the dashed lines corresponding to tight linkage (eq. (S20)) and the solid lines to loose linkage (eq. (4)). The right panel corresponds to the recessive model. For the recessive model, a strong barrier cannot be form if the loci are in tight linkage (no dashed lines) and the solid lines are obtained from numerical solution of the evolution equations. In both panels, the squares correspond to results for the equivalent discrete-time model that allows for strong selection, assuming that *A* and *B* are on different chromosomes.

For the codominant model in loose linkage, we proved that a neutral continental adaptation, *β* = 0, is the easiest condition to form a barrier that exceeds 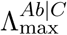; by easiest condition we mean that it requires the least amount of negative epistasis, as it maximizes equation (4). A neutral *B* allele does not contribute to the maximum amount of local adaptation, therefore all local adaptation can be captured by the *A* adaptation 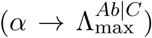. At the same time, if *B* is not advantageous on the island, direct selection is not acting against the maintenance of the DMI and does not reduce the strength of genetic barrier.

For the codominant model, having *A* and *B* in tight linkage requires less epistasis to form a genetic barrier that exceeds 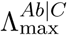 than in loose linkage. That is because selection against hybrids is expressed for both linkage architectures, but the migration cost is paid only once if *A* and *B* are in tight linkage, but twice if in loose linkage. Therefore, it is easier to form a strong barrier in tight linkage. For the recessive model, 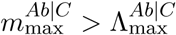 is possible only if there is recombination between the two loci, otherwise the incompatibilities are never expressed and selection against hybrids is inactive.

The discrete-time model is qualitatively similar to the continuous-time model as illustrated in Figure 4(a): The different dots correspond to the minimal conditions on the strength of epistasis to observe a genetic barrier stronger than the maximum amount of local adaptation in the discrete-time codominant model. The fact that both the continuous and discrete time model are qualitatively similar is crucial, since the formation strong barriers require strong epistatic interactions, for which the equivalence between continuous and discrete-time model is no longer ensured.

## Discussion

We here show that interactions between three loci can be sufficient to confer strong reproductive isolation between two populations in parapatry, and the evolution of this barrier is possible in the presence of ongoing gene flow. We first establish that in the absence of epistasis or under a large number of “simple” epistasis schemes (as described above), the amount of local adaptation between well-adapted types in both populations is a hard limit for the strength of a genetic barrier. We then describe a simple 3-locus scenario in which a much stronger barrier can evolve. Crucially, the scenario relies on cryptic epistasis, i.e., epistasis between the divergent alleles and a derived background allele that fixes in both populations. In this case, a strong barrier is possible if a classical 2-locus DMI is stabilized by positive epistasis of both interacting partners with such a background allele. Since the strength of the genetic barrier relies on strong selection against hybrids, this phenomenon requires sufficiently strong recombination between the interacting loci to be observable in haploid populations. In diploids, where hybrid genotypes also form without recombination, codominance of the incompatibilities and tight linkage between the loci involved in the initial DMI provide the best conditions for the evolution of strong reproductive isolation.

### Postzygotic reproductive isolation and ecological speciation

The accumulation of genetic incompatibilities due to selection or drift is a standard mechanism to explain the evolution of reproductive isolation between two allopatric populations [2]. In the presence of gene flow, however, each new incompatible mutation faces a fitness deficit. Theoretically, a contribution to local adaptation by each of these mutations can make up for this deficit. Indeed, it has been shown that the accumulation of locally adaptive mutations between two parapatric populations can result in genetic barriers to gene flow of arbitrary strength [11, 9]. Realistically, however, the maximum amount of local adaptation that is available (as a function of the differences in the external environment) between two populations will often be limited: while migrants from nearby habitats often have a fitness deficit relative to locals, they are usually not entirely lethal or infertile. Imposing such an upper bound immediately renders an upper bound for the strength of a genetic barrier. In the presence of epistasis and genetic incompatibilities, fitness deficits of hybrids may be much larger than the ones of migrants, opening up the potential for a stronger barrier. Nevertheless, our results show that for most models with simple epistasis, local adaptation is still a limit for the amount of gene flow that a barrier, built in parapatry, can sustain: *m*_max_ ≤ Λ_max_. This limit holds 1) for all 1 and 2-locus models, 2) for all models in which all loci are tightly linked, 3) for models with only island adaptations or deleterious continental mutations, and 4) for models with only negative epistasis between continental and island mutations.

### Cryptic epistasis enables the formation of a genetic barrier stronger than the maximum amount of local adaptation

Conceptually, speciation in the presence of gene flow requires a fitness landscape in which (at least) two peaks are connected via a high-fitness ridge of single-step mutations. Yet, to exceed the limit imposed by the maximum amount of local adaptation, any recombinants between the peak genotypes have to be strongly deleterious. This can be achieved by what we term “cryptic” epistasis, i.e, when the interaction with (at least) a third derived allele turns the high-fitness ridge that allowed for the evolution of an initial DMI into a fitness valley. Importantly, this third allele must fix in the population, or otherwise the high-fitness ridge is not yet fully interrupted.

In a minimal model, three loci are necessary to form the required underlying fitness landscape. In this landscape, the first mutational step corresponds to the establishment of initial differentiation between the two populations, which requires some local adaptation (either on the continent or on the island) at the respective locus. The second mutation generates a derived-derived incompatibility with the first adaptation (for the equivalence with other types, see [5]). At this point, two fitness peaks correspond to the two derived haplotypes, one of which is fixed on the continent, whereas the other dominates the island. These peaks are still connected via a high-fitness recombinant, namely the ancestral haplotype, which is always segregating due to migration and recombination of the two derived haplotypes. Finally, a third mutation occurs on the continent; this adaptation is deleterious in the background of the ancestral haplotype, but advantageous in the presence of both previous mutations. If this third mutation fixes on both the continent and the island, recombinants between the dominant haplotypes on the continent and the island (each of which inhabit a fitness peak) are always deleterious. As a consequence, the resident island genotype can now withstand much stronger gene flow than suggested by the fitness differences between the two derived haplotypes.

For a hypothetical example of cryptic epistasis, assume that mutations at loci *A* and *B* correspond to adaptations leading to specialization for the prevalent food source on island and continent, respectively. Both come with a (large) cost of catching/exploiting the other one, such that *AB* individuals are not good catching/exploiting either. Mutation *C* makes individuals stick to a single foraging pattern, which is bad for the *ab* generalists, but good for both specialists, and may thus fix in both populations.

DMIs have been investigated mainly with respect to negative pairwise epistasis [17, 18, 5, 19, 13]. Here, we showed that more complex epistasis can significantly alter the potential for the evolution of reproductive isolation in parapatry. A key player in our minimal model of strong reproductive isolation is an allele that becomes fixed across both diverging populations during the course of the speciation process. The possibility that globally fixed mutations are involved in the speciation process complicates the challenge of inferring speciation genes and reconstructing the evolutionary trajectory of the incipient species. Specifically, these fixed mutations, responsible for what we term cryptic epistasis, will only be detected as divergent with a sister-clade and they will not appear in F1 and F2 hybrid viability analysis [20, 21], thus their role in the speciation process may easily be overlooked.

The importance of complex (non-negative pairwise) epistatic interactions in speciation has been stressed in several studies. Fraisse *et al*. [22] compiled a list of studies with DMIs of higher order than pairwise interactions and using the framework of Fisher’s geometric model, showed that complex DMIs are likely to play an important role in the speciation process. In a model of secondary contact [23], divergent gene clusters with complex incompatibilities, but without any local adaptation (neutral gene networks), can be maintained in the face of secondary gene flow. The less connected the neural network is, the easier it is to maintain the divergence. Since all steps on the network are neutral, however, divergence can never evolve in the presence of gene flow and an allopatric phase is always necessary.

### Scope and limits of our model

The results presented here were derived using an analytical framework, complemented with some numerical calculations. To do so, we used a continuous time approximation, which has the disadvantage of having parameters that are meaningful only in relationship with each other. We confirmed that we observe a qualitatively similar pattern in a discrete time scenario, where parameters can be transposed to natural cases. Furthermore, we investigate this question under an infinite population size model. Adding genetic drift to the model is of great interest as temporal dynamics, as well as drift, may impact the final outcome. Adding drift may probably weaken the genetic barrier since the island population will be smaller. However, it may favor the introgression of background mutations from the continent to the island and therefore accelerate the formation of strong genetic barriers. Similarly, we focus mainly on cases of linkage equilibrium. Feder et al. [24] showed that strong linkage disequilibrium between many loci may trigger a genome-wide reduction in gene flow, “genome congealing” (sensu Turner [25, 26, 27]). It will be interesting to see how these two mechanisms may combine during the speciation process. Finally, we only observed the evolution of these strong genetic barriers when the *C* mutation fixed on the island, but could not exclude the possibility that strong barriers can evolve even if the *C* allele remains polymorphic on the island.

In our minimal model, a lot of deleterious hybrids will be generated which comes at a cost for the island population. Co-existence of the “island species” and the “continental species” in this case thus relies on a sufficiently large population size on the island, such that the “island types” are always in the majority relative to the continental migrants (Fig. S8). In this case, the continental migrants suffer more from matings with the island types (since continental types will mainly produce inviable hybrid offspring). The dynamics may change if subsequent evolution of prezygotic isolation strengthens the genetic barrier without requiring any further local adaptation. Indeed, our model should provide a favorable scenario for such reinforcement [28, 29, 30]. However, even if all types avoid matings with the opposite type, the continental type may eventually still swamp the island due to migration pressure. This would depend on the details of the assortment mechanism and may be precluded if mate choice comes at a cost.

### A route to parapatric speciation?

Hybrid incompatibilities have been proposed as an engine of speciation in allopatry, where simple accumulation of individually neutral but negatively interacting mutations “almost necessarily” leads to a “snowball effect” and eventual reproductive isolation [17], a process which is impeded in the presence of any amount of gene flow [5]. In a similar vein, the accumulation of locally adapted alleles was proposed as a natural engine of speciation in parapatry [11]. By studying the interaction of local adaptation and hybrid incompatibilities in the presence of gene flow, our previous [5][9] and current work challenges the view of parapatric speciation as a gradual and monotonous process that is mainly driven by local adaptation.

We have previously shown that some local adaptation is indeed a necessary ingredient for the evolution of a genetic barrier in the presence of gene flow [5], and that this barrier can either grow or shrink as additional mutations appear [9]. Here, we show that in a large class of models with simple fitness landscapes, the ecological differentiation is an upper bound for the strength of a genetic barrier that can evolve in the presence of gene flow. Thus, if local adaptation is limited (which it realistically is), also the potential for the evolution of reproductive isolation in parapatry is usually limited.

Importantly, we also discovered specific fitness landscapes that combine locally adapted alleles with specific epistatic interactions, which enable the evolution of much stronger genetic barriers and even complete isolation in the presence of gene flow. Whether strong reproductive isolation between parapatric populations might indeed evolve through the combination of local adaptation and epistasis described here is thus dependent on the existence of the necessary fitness landscapes in nature. If they exist, the route to strong reproductive isolation could require only a small number of mutational steps. If such fitness landscapes do not exist, strong postzygotic reproductive isolation in the presence of gene flow may never be reached even after a very long time. An important conclusion from our work is thus a strong dependence of the feasibility of parapatric speciation on the underlying genetics, which makes it difficult to infer and predict.

## Acknowledgments

We thank R. Bürger, M. Servedio, C. Vogl, S. Mousset, I. Fragata, I. Höllinger and the Biomathematics Group at the University of Vienna for helpful discussion and comments on the manuscript, and the editor and the two anonymous reviewers for their valuable suggestions that have improved this manuscript. A.B. was supported by the Marie Curie Initial Training Network INTERCROSSING. C.B. is grateful for support by EMBO Installation Grant IG4152. A.B. and C.B. were supported by ERC Starting Grant 804569 - FIT2GO.

